# Rapid quantification of polyhydroxyalkanoates accumulated in living cells based on green fluorescence protein labeled phasin: The qPHA method

**DOI:** 10.1101/2022.03.02.482659

**Authors:** Xu Liu, Dianjie Li, Xu Yan, Zonghao Zhang, Shuang Zheng, Jingpeng Zhang i, Fuqing Wu, Fangting Li, Guo-Qiang Chen

## Abstract

Polyhydroxyalkanoates (PHA), are microbial polyesters with possibility to replace non-biodegradable petro-plastics. No rapid *in situ* PHA quantitation method has been available for the past 40 years to replace the traditional method which is complicated, time and labor consuming. Quantification of PHA in living cells were finally developed from fluorescence intensities generated from green fluorescence protein (GFP) fused with the *Halomonas bluephagenesis* phasin proteins attached on the PHA granules. Phasins PhaP1 and PhaP2 were used to fuse with GFP which reflects PHA accumulation with an R-square over 0.9, respectively. Also, a standard correlation was established to calculate PHA contents based on the fluorescence and cell density recorded via a microplate reader with R-square over 0.95 when grown on various substrates, respectively. The PhaP2-GFP containing *H. bluephagenesis* was applied successfully to quantify PHA synthesis in a 7.5 L fermenter with high precision. The method is named qPHA.

## 1. Introduction

Recently, global plastics production reached approximately 370 million tons annually with estimation of 79% accumulated in landfill or natural environments as unrecyclable and non-biodegradable plastic wastes that pollute our environments and threaten wild lives (Geyer et al., 2017; PlasticsEurope, 2020). As a result, biodegradable plastics have flourished in the last decades including polylactide (PLA) and polyhydroxyalkanoates (PHA) synthesized by many bacteria as carbon and energy storage compounds (Chen, 2009). PHA consist of various hydroxyalkanoate monomers leading to various material properties (Chen, 2009; Tan et al., 2021). PHA can be manufactured as daily plastics, textiles, medical implants, drug delivery carriers and other useful products (Chen, 2009; Tan et al., 2021). Next generation industrial biotechnology (NGIB) based on extremophilic bacteria has been developed to overcome the economic and technological challenges involved in PHA commercialization. *Halomonas bluephagenesis* TD01, short as *H. bluephagenesis*, a halophilic bacterium isolated from a salt lake in Xinjiang Province of China, has been developed as a chassis for NGIB for its non-sterile fermentation strategy under high-salt and alkali culture condition (Chen & Jiang, 2018; Tan et al., 2011).

Gas chromatographic (GC) analysis is the common approach to quantify the intracellular PHA. To perform the GC study, the microbial cells containing intracellular PHA should first be harvested via centrifugation, washing and then lyophilized. Subsequently, the weighted cell or PHA samples are heated to 100℃for 4 hours to depolymerize PHA under acidic condition in the presence of methanol and chloroform, so as to convert PHA monomers to methyl-hydroxyalkanoates for GC analysis (Braunegg et al., 1978). The entire PHA analysis process lasts nearly two days, preventing the *in situ* process control based on PHA percentage accumulation during the fermentation processes. Although gas chromatography is the classic method to determine PHA content and polymer compositions, its labor and time-consuming process also generates harmful methanol and chlorinated solvents as waste solvents (Braunegg et al., 1978). Although some staining methods were used for PHA detections, including Nile red, BODIPY, double-staining with BODIPY 493/503-SYTO 62 and other lipophilic fluorescent dye (Karmann et al., 2016; Kettner & Griehl, 2020; Lee et al., 2013; Zuriani et al., 2013), these stainings methods could not meet the changing fermentation needs because their measurements were not instantaneous. To perform the staining methods, the cells should be stained and washed several times and the results of the stainings were easily influenced by the operator skills.

PHA are stored by bacteria as water-insoluble inclusion granules enveloped by granule-associated protein phasins (Dennis et al., 2008; Pieper-Furst et al., 1995), that improve the bacterial stress resistance abilities and contribute to the enhanced PHA accumulation (de Almeida et al., 2011; de Almeida et al., 2007; Ushimaru et al., 2015). Furthermore, phasins have been developed for various applications including drug delivery systems, protein purification and bio-surfactants (Wang et al., 2008; Wei et al., 2011; Yao et al., 2008). Previous studies indicated that the amount of PHA accumulation was related to the expression level of phasin in many strains, such as *Cupriavidus necator* (*C. necator,* formerly designated *Ralstonia eutropha* and *Alcaligenes eutropha*) and *Paracoccus denitrificans (P. denitrificans*) (Maehara et al., 1999; Potter et al., 2002; Velazquez-Sanchez et al., 2020; York et al., 2001a).

This study aimed to develop a rapid and *in situ* method based on relationship of phasin and PHA accumulation to quantify the intracellular PHA percentage content in living *H. bluephagenesis,* so as to monitor and then control the PHA production processes.

## 2. Materials and Methods

### 2.1 Bacterial strains, plasmids and growth conditions

The strains and plasmids used in this study are listed in Tables S1. *Halomonas bluephagenesis* TD01 (NCBI number: txid2778948) and its mutant strains were cultured in a LB60 medium containing 60 g/L NaCl as seed inoculant. *Escherichia coli* S17-1 cultured in LB medium was used as the host for plasmid construction and the conjugation donor. Plasmid was constructed by Gibson Assembly method. The DNA fragments were amplified by PCR using Q5® High-Fidelity DNA polymerase (New England Biolabs Inc., USA).

To prepare the medium, LB medium was mixed by 10 g/L tryptone (Analytical reagent, Oxoid, England), 5 g/L yeast extract (Analytical reagent, Oxoid, England) and 10 g/L NaCl (Analytical reagent, Sinopharm Chemical Reagent Co.,Ltd., China). The NaCl concentration was changed to 60 g/L in the LB60 medium. For shake-flask PHA production, a defined minimal medium MM60 containing 6% NaCl, 0.02% MgSO4, 1.0% Na2HPO4·12H2O, 0.15% KH2PO4, 1.0% trace element solution I, 0.1% trace element solution II, 0.1% yeast extract and 3% glucose was used. The urea concentration was prepared depended on the specific needs. Chloramphenicol (25 μg mL^−1^), kanamycin (50 μg mL^−1^), and spectinomycin (100 μg mL^−1^), are purchased from BioDee Biotechnology Co., Ltd. (Beijing, China), they were added to the above media whenever necessary. All the culture and fermentation experiments were carried on at 37°C. And all chemicals were obtained from Shanghai Sangon Biotech Co., Ltd.

### 2.2 Fluorescin fusion in genome using CRISPR/Cas9 and conjugation

The GFP was fused with the PhaPs protein using a linker SGSS *in situ* by CRISPR/Cas9 in *Halomonas bluephagenesis* (Qin et al., 2018). Three single-guide RNAs (sgRNA) were synthesized from the downstream of three *phaPs* and the homologous arms were designed to delete the stop codon (TAA) of *phaPs*. The sequence of linker (amino acid sequence SGSS) and *gfp* were constructed in the donor plasmids with the homologous arms designed previously. And, the *H. bluephagenesis* or its mutant could be transformed with pQ08 plasmid which expresses Cas9 protein via conjugation. Selected under chloramphenicol, the donor plasmid was transformed into the *H. bluephagenesis* harboring the pQ08 plasmid. Then, colony PCR and DNA sequencing were conducted to verify the correctly edited strain grown in LB60 plate containing chloramphenicol and spectinomycin. Subsequently, both donor and Cas9 plasmids forming the CRISPR/Cas9 system were cured from *H. bluephagenesis*, so that the genome edited strains became plasmid-free.

Conjugation was an efficient method to transform plasmids into *H. bluephagenesis*. The *E. coli* S17-1 harboring the plasmid and the target *H. bluephagenesis* strain were cultured for approximately 4 h to reach an OD600 of 0.3 to 0.5. the cells were harvested at ratio 1:1 via centrifugation at 4000 g for 2 min, washed twice using a fresh LB medium and finally mixed together. After resuspended with 50 μL fresh LB medium, the mixture were cultured on antibiotic-free LB20 plates at 37°C for 6 h. Finally, the bacterial lawn was directly spread onto a LB60 agar plate containing appropriate antibiotics followed by incubation at 37°C for 48 h.

### 2.3 Growth dynamic studies in shake flasks

The shake flask studies began with streaking glycerol stored cells in fresh LB60 agar plates to obtain single colonies. After selecting right single colonies from the plates, the cells were transformed to LB60 medium and incubated at 200 rpm overnight to acquire the first seed culture. Subsequently, the second seed culture at OD600 around 1.0 was acquired from 1% inoculation in a 20 mL fresh LB60 medium for 8 h under the same culture condition. The fermentation medium was based on 50 mL MM60 medium mentioned previously using the 5% second seed culture in a 500 mL conical flask at 200 rpm. For the dynamic growth studies, a 5 mL sample was taken at intervals from the shake flask under sterile conditions over the growth period.

### 2.4 Growth and PHA production in 7.5 L bioreactor

The seed culture was prepared in the same way as the shake flask ones. A 300 mL second seed culture was inoculated into the fermenter containing 3L MM60 culture medium supplemented with 5 g/L yeast extract and 2 g/L urea at the beginning. While the feeding solution I and II contain 2 g/L urea and 800 g/L glucose in 0.4 L volume, and only 800 g/L glucose, respectively. The flow rate of feeding the solutions was controlled to maintain around 10 g/L glucose in the bioreactor during the cultivation. 35 mL sample was acquired each sampling from the bioreactor for analyzing the cell dry weights (CDW) and PHA content when necessary.

### 2.5 PHA quantification via gas chromatography (GC)

The samples obtained from the shake flasks were harvested via centrifugation and washed using distilled H_2_O twice, followed by freeze-drying overnight in a lyophilizer. The so generated CDW was incubated at 100°C for 4 h in 2 mL chloroform and 2 mL esterification fluid containing 3% concentrated 98% sulfuric acid and 0.1% benzoic acid in methanol. Finally, the clear chloroform phase containing hydroxyalkanoate methyl esters was used for gas chromatography analysis using a GC-2014 (Shimadzu, Japan) after 1 mL water extraction. P3HB (99.9%, Sigma-Aldrich, Germany) and γ-butyrolactone (99.9%; Shanghai Macklin Biochemical Co., Ltd., China) were used as the respective standards. The band representing benzoic acid was regarded as the internal standard.

### 2.6 Flow cytometry analysis

The samples were diluted in proportion of 0.5% with PBS and 10 μL of the diluted solution were sampled in every measurement using a flow cytometer (LSRFortessa4, BD, USA). GFP were excited at 488 nm, the signals of fluorescence intensity were captured on an FITC (voltage at 420 V), FSC (forward scatter, voltage at 450 V), and SSC (side scatter, voltage at 185 V) channel at a rate of 0.5 μL s^−1^ for 20 s. Their means of all tested cells (10,000) were recorded as FCM-FI.

### 2.7 OD_600_ and fluorescence intensity recorded using a microplate reader

The samples were diluted to an OD600 between 0.3 to 0.8 using ultra-pure water. 200 μL diluted bacterial solution of each sample was added into a 96-wells microplate (Corning, USA) to measured absorbance at 600 nm and fluorescence intensity with excitation/emission wavelength at 488/518 nm using a microplate reader (VarioskanFlash, Thermo Scientific, USA). The diluted bacterial solution of each sample was also centrifuged. Subsequently, the supernatant was measured for normalization.

### 2.8 OD_600_ and fluorescence intensity recorded using a visible spectrophotometer and a fluoro-spectrophotometer

The samples were diluted to OD600 between 0.3 to 0.8 using ultrapure water in volumetric flasks. 3 mL diluted bacterial culture of each sample was added into quartz cuvette to measure absorbance at 600 nm via visible spectrophotometer (V-5600, Shanghai METASH, China), and fluorescence intensity with excitation/emission wavelength at 470/514 nm via fluoro-spectrophotometer (F97pro, Shanghai Lengguang Technology, China).

### 2.9 Observation under Structure Illumination Microscopy (SIM)

The strains with GFP labeled PhaP were observed after 48 hours fermentation in MM60 medium (1 g/L urea) via shake flask experiment under N-SIM super resolution microscope (Nikon, Japan).

The equipment was the N-SIM microscope with the SR Apo TIRF 100X oli objective. And the illuminator was the N-SIM illuminator. The filter settings were 488 nm for the GFP observation and the 561 nm for the Nile Red staining. The excitation wave length were 488 nm and 561 nm, respectively. The SIM method was the 3D SIM. And the reconstruction parameters are as set as described below.

488 nm (Illumination Modulation Contrast: 1.52, High Resolution Noise Suppression: 1.40, Out of Focus Blur Suppression: 0.28)

561 nm (Illumination Modulation Contrast: 1.52, High Resolution Noise Suppression: 2.28, Out of Focus Blur Suppression: 0.20)

And for the Nile Red staining, the straining method was derived from the reference (Lee et al., 2013).

### 2.10 Data analysis and fitting of the relationship between the PHA content and the single cell fluorescence intensity

To assay relations of the FCM-FI (f) and the PHA content (PHA), their average values and standard deviations were calculated at the same time. The average data were fitted with the linear function Eq. (3) and the exponential function described below.

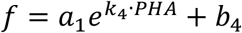

Here, *f* is the single cell fluorescence intensity and *PHA* is the PHA content. We used the standard nonlinear least squares fitting tool in Python, and the R-square is defined as,

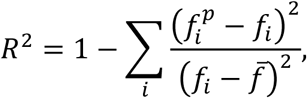

where *f_i_* is the experimental single cell fluorescence intensity, 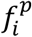 is the predicted value by the fitting functions, and 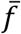 is the average intensity.

### 2.11 Establishment of relationship among OD_600_, GFP fluorescence and the PHA content

The diluted OD_600_ was multiplied via the dilution ratio to gain a new OD600 (O) for following data fitting. For instance, if 20 μl sample is diluted to 200 μl into the microplate reader and the corresponding OD_600_=0.5, the modified OD_600_=0.5/(20/200)=5. Same modification was applied on the GFP intensity to obtain a new GFP fluorescence intensity (F) for data fitting.

We directly fitted OD_600_, FI and the PHA content (PHA, P) with the expression Eq. (4) by the standard nonlinear least squares fitting tool in Python. The R-square is defined by the PHA content,

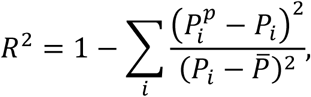

where *p_i_* is the experimental PHA content, 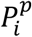 is the predicted PHA content, and 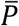 is the average experimental PHA content.

When fitting the data of IPTG induced experiments and bioreactor experiments, the standard parameter values (*p_i_*) were used as the initial values 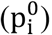, and the bounds were set between 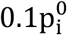 and 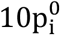. During the parameter estimation of IPTG data, the instrument related parameters b1 and b2 were fixed as standard values. For the bioreactor experiments, the parameters were changed accordingly.

### 2.12 Dynamic imaging of single cells

The cells were cultured in 4 mL LB60 medium for 12 hours, followed by sampling 250 μL aliquots, then diluted with 750 μL fresh LB60 medium for imaging. The diluted cells were injected to the microfluidic chips, and continuously injected with MM60 (1 g/L urea, 30 g/L glucose) as the medium. The flow rate was set to 50 μL h^−1^. The cells were precultured in microfluidic chips for 4 hours before imaging. The images were captured for 6 hours under the fluorescence microscope (TiE, Nikon, Japan), where the time interval was 5 minutes and the temperature was 37℃. The movie was filmed based on the images by ImageJ (Movie 1).

## 3. Results and Discussion

### 3.1 Construction of *Halomonas bluephagenesis* strains with GFP labeled phasin

The PHA producing *H. bluephagenesis* contains three phasin genes including *phaP1*, *phaP2* and *phaP3* (Shen et al., 2019). The CRISPR/Cas9 approach was used to construct the PhaP-GFP fusion protein using a linker. Several studies suggested that the PhaP linked with labeled fluorescent protein in its N- or C-terminal could both bind to the PHA granule. And these results were determined by microscopic studies and immunoblotting (Hauf et al., 2015; Neumann et al., 2008). PhaPs in our study were chosen for its C-terminal fusion. As described in the Materials and Methods, the GFP labeled strains were successfully constructed containing the PhaPs and GFP fusion via a linker. Finally, three single-*phaP*-labeled strains based on wildtype *H. bluephagenesis* were obtained, namely *H. bluephagenesis* strains TDP1-GFP, TDP2-GFP, TDP3-GFP (Tables S1).

In our previous study, the *phaP1*-deleted *H. bluephagenesis* TDΔ*phaP1* can produce larger PHA granules than the wildtype does, this is very convenient for easy downstream PHA recovery and purification processes via centrifugations (Shen et al., 2019). Thus, the other two single-*phaP*-labeled strains of *H. bluephagenesis* based on TDΔ*phaP1* were constructed, namely *H. bluephagenesis* TD*ΔphaP1*P2-GFP and TDΔ*phaP1*P3-GFP (Tables S1).

All GFP labeled *H. bluephagenesis* strains, and the PHA synthase (PhaC) deleted strain *H. bluephagenesis* TDΔ*phaC*P1-GFP (which could not accumulate PHA) with GFP labeled PhaP1, were observed after 48 h shake flasks experiment, respectively, under Nikon structured illumination microscopy (N-SIM), a super resolution microscope (Fig. 1). When the strain could synthesize PHA (like TDP1-GFP group), some green circles were observed because PHA granules were surrounded by GFP-labeled PhaP protein. Compared between the TDP1-GFP and the TDΔ*phaC*P1-GFP groups, when the strain could not synthesize PHA, the PhaP-GFP fluorescence intensity was weak and could not form green circles. For the PhaP1 is the most abundant phasin in *H. bluephagenesis* (Shen et al., 2019) PhaP2 and PhaP3 were able to form the complete green circles only in the *phaP1*-defient strains (TD*ΔphaP1*P2-GFP and TDΔ*phaP1*P3-GFP, respectively). To further confirm the relationship between the green circles formed by PhaP-GFP and the PHA granules, the Nile Red staining were applied in the labeled strains. The co-localization results were shown in the Supplementary material. The green PhaP-GFP was tend to encircle the red PHA granules stained by Nile Red.

**Figure. 1.**
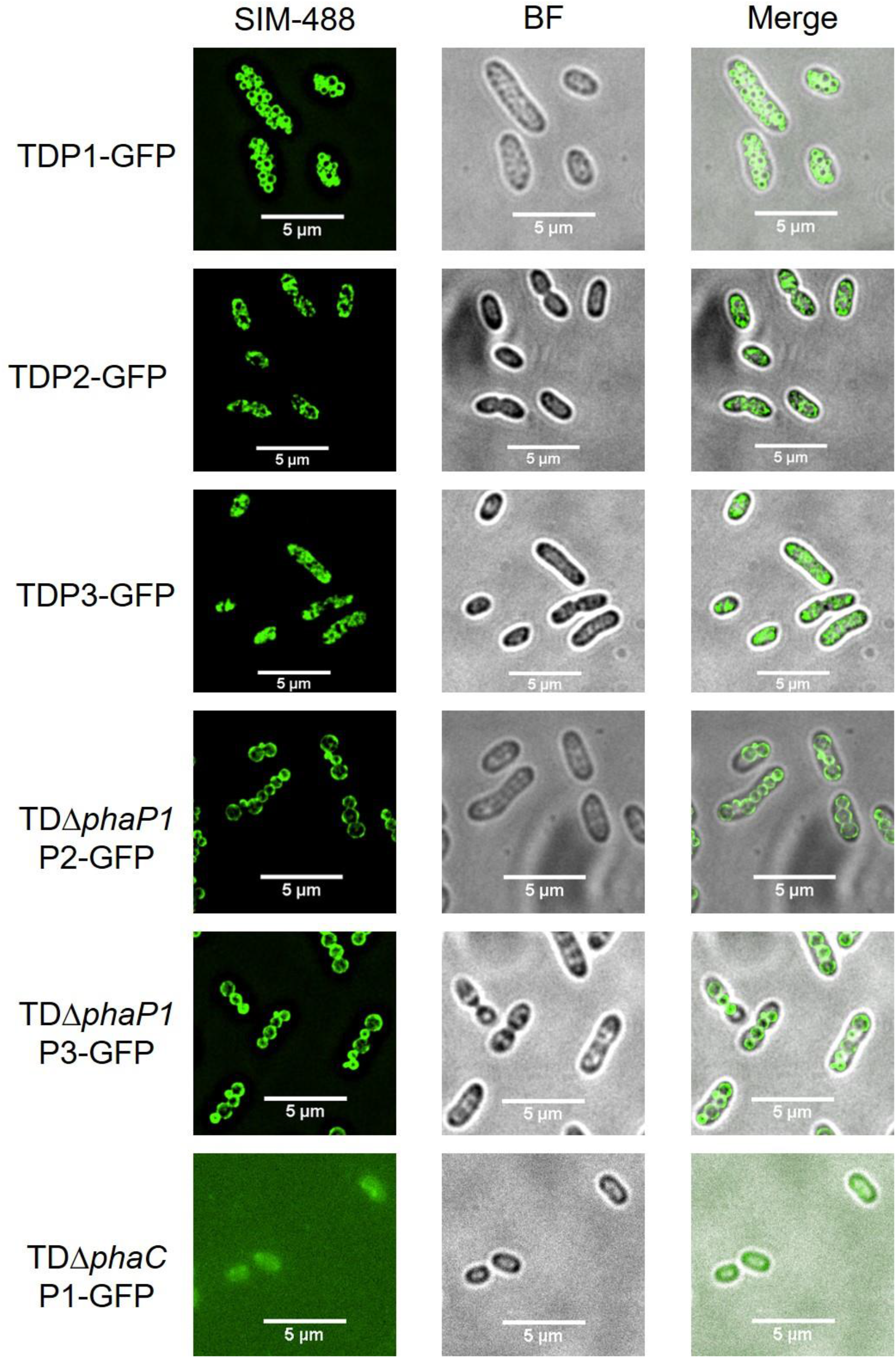
Design of the PHA quantitation method (qPHA) based on GFP labeled PhaPs. Microscopic observation of the GFP labelled PhaPs in *H. bluephagenesis.* The first, fourth, fifth lines showed the green circles in TDP1-GFP, TDΔ*phaP1*P2-GFP and TDΔ*phaP1*PhaP3-GFP, respectively. the second and the third lines showed the uncompleted green circles in TDP2-GFP and TDP3-GFP, respectively. The last line showed the PhaP1-GFP does not form circles in *phaC* deleted *H. bluephagenesis* TDΔ*phaC*P1-GFP. All the pictures of strains were shown in three columns, including SIM-488 channel, bright filed channel and merged channel. Samples were observed after 48 h in shake flasks.

All the *phaP* labeled strains were captured under microfluidics to observe the process of PHA accumulation and the Movie 1 was an example of *H. bluephagenesis* TDΔ*phaP1*P2-GFP. During the microfluidics culture, the fluorescent intensity of each cell was increasing obviously.

### 3.2 Relationship of fluorescence intensities with PHA contents

Flow cytometry was used to study the fluorescence intensities (FCM-FI) of the six GFP labeled PhaP strains including *H. bluephagenesis* strains TDP1-GFP, TDP2-GFP, TDP3-GFP, TDΔ*phaP1*P2-GFP, TDΔ*phaP1*P3-GFP and TDΔ*phaC*P1-GFP grown in shake flasks, respectively. As described in Fig. 2a, the PHA content data were obtained through the GC method (including centrifugation, lyophilization, esterification and gas chromatography analysis steps) and the FCM-FI was measured using the sample acquired from the same time. During the shake flask experiment, the *phaC*-deficient strain could not accumulate PHA and the FCM-FI was also extremely low. The FCM-FI of each cell were observed in an approximately positive correlation pattern to increase with enhanced PHA accumulation in other GFP-labeled cells grown in the presence of high nitrogen concentration condition (2 g/L urea) (see Supplementary material), even though various PhaPs labeled GFP emitted various FCM-FI, respectively. However, the positive correlation pattern of the FCM-FI and PHA content in the above five strains were disrupted under the low nitrogen concentration condition (0.5 g/L urea) (see Supplementary material), as the cells cannot synthesize enough GFP under N deficiency. These results indicated that nitrogen supply influences the relationship between FCM-FI and PHA content. The closely linear relationships of the FCM-FI and PHA content could be exploited to quantify the intracellular PHA content under specific nitrogen conditions. To understand how nitrogen (urea) concentration affects relationships of the FCM-FI and PHA content, *H. bluephagenesis* TDP1-GFP was grown in the presence of 0.8 g/L to 1.8 g/L urea with 0.2 g/L interval in the MM60 medium in shake flasks. Results showed that the FCM-FI increased with the PHA accumulation above the presence of 1 g/L urea (Fig. 2b). To further refine the relationship between the FCM-FI and the PHA content, the linear and exponential functions were chosen to fit the experiment data with the FCM-FI under various urea concentrations, respectively (Fig. 2c). Simulation curves obtained from two types of functions fitted very well with the experimental data with a R-square around 0.90 except for that of the 0.8 g/L urea concentration (Fig. 2d). Otherwise, 1 g/L urea concentration group reach the highest PHA content in the groups which had the positive correlation between the FCM-FI and the PHA content. Based on the above results, 1 g/L of urea was chosen for following experiments to satisfy the opposite demands for both PHA production and the positive correlation pattern.

**Figure. 2.**
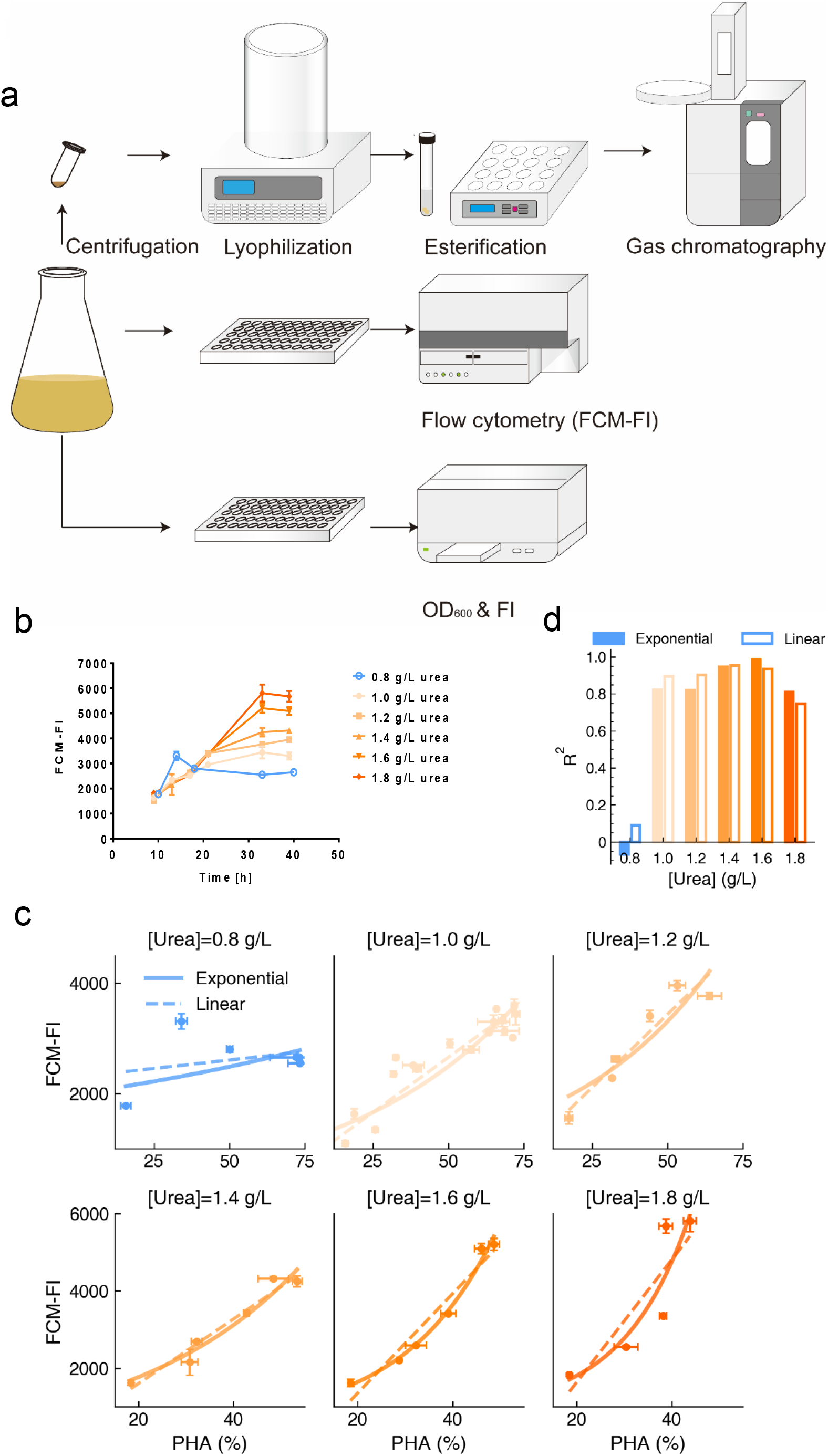
Fluorescence intensities generated by. H. bluephagenesis TDP1-GFP grown at different urea concentrations, respectively. (a) Schematic illustration of the qPHA method and traditional GC method. The GC method comprises four steps (centrifugation, lyophilization, esterification and gas chromatography analysis). The qPHA method only comprises one step measurement via flow cytometer or microplate reader. (**b**) Dynamic fluorescence intensities of the cells grown at different urea concentrations in shake flasks, respectively, were measured using flow cytometry (FCM-FI). (**c**) Exponential and the linear functions were used to fit the relationship between the FCM-FI and the PHA percentage content at different urea concentrations, respectively. (**d**) The R-square values of **Fig. 2c** were shown as the columns. The FCM-FI was calculated by the mean of 10000 cells. The solid and hollow columns represented exponential functions and linear functions, respectively. All data represent the mean of n = 3 biologically independent samples and error bars show s.d.

In the same time, the growth was analyzed in the labeled strains under 1 g/L urea. After 48 hours shake flask experiments, the cell dry weights (CDW) and PHA contents of the labeled strains (TDP1-GFP, TDP2-GFP, TDP3-GFP, TDΔphaP1P2-GFP and TDΔphaP1P3-GFP) and the control strains (TD01 and TDΔ*phaP1*) were measured by lyophilization and gas chromatography analysis (see Supplementary material). There seems no difference between the labeled strains and the control strains. These results suggested that the GFP label did not influence the strain growth and PHA accumulation. Moreover, The PHA production was compared between the 0.5 g/L urea group and the 1.0 g/L urea group in TD01 strain. The 0.5 g/L urea concentration was our normal shake flask culture condition (Tan et al., 2011). The shake flake results showed that the 1.0 g/L group got higher PHA yield. Although it had lower PHA content but higher CDW (see Supplementary material).

### 3.3 Selection of suitable PhaP for GFP fusion to fit FCM-FI to PHA contents

*H. bluephagenesis* contains three phasins including PhaP1, PhaP2 and PhaP3 that are all fused with GFP for FCM-FI studies, respectively. Different PhaPs were reported to have specific regulatory functions in *C. necator* (Neumann et al., 2008; Potter et al., 2005). To further select the most suitable PhaP among the three phasins for characterization of intracellular PHA contents, *H. bluephagenesis* strains TDP1-GFP, TDP2-GFP and TDP3-GFP were studied taking the GFP-labeled PhaC containing *H. bluephagenesis* TDC-GFP (TDC-GFP) as a negative control grown all in the presence of 1 g/L urea. PhaC was the PHA synthase, which was also located in PHA granule surface and could catalyze the hydroxyacyl-CoA into polyhydroxyalkanoates (Peters & Rehm, 2005).

*H. bluephagenesis* TDP1-GFP and TDP2-GFP showed a much close relationship between FCM-FI and PHA contents compared to that of *H. bluephagenesis* TDP3-GFP and TDC-GFP grown all in 1 g/L urea containing MM60 in shake flasks, respectively (Figs. 3a-3d). When expressed using exponential and linear functions, the relationships of FCM-FI and PHA contents in all the different GFP-labeled proteins containing *H. bluephagenesis* strains were highly similar or even overlapped, with FCM-FI of *H. bluephagenesis* TDP1-GFP and TDP2-GFP fitting much better to PHA contents (Figs. 3e and 3f) than that of *H. bluephagenesis* TDP3-GFP and TDC-GFP (Figs. 3g and 3h). The R-square of exponential and linear functions used to fit the experiment data demonstrate that PhaP1 and PhaP2 are more suitable than PhaP3 to reflect PHA contents as both their R-squares of exponential and linear functions reach to 0.90 compared to less than 0.8 from PhaP3(Fig. 3i). And the R-square of the negative control TDC-GFP was only 0.3, which confirm the unique coordination relationship between PhaP protein and PHA contents. These results firmly suggest that PhaP1 and PhaP2 are more suitable as the target GFP labeled protein for quantification of intracellular PHA contents.

**Figure. 3.**
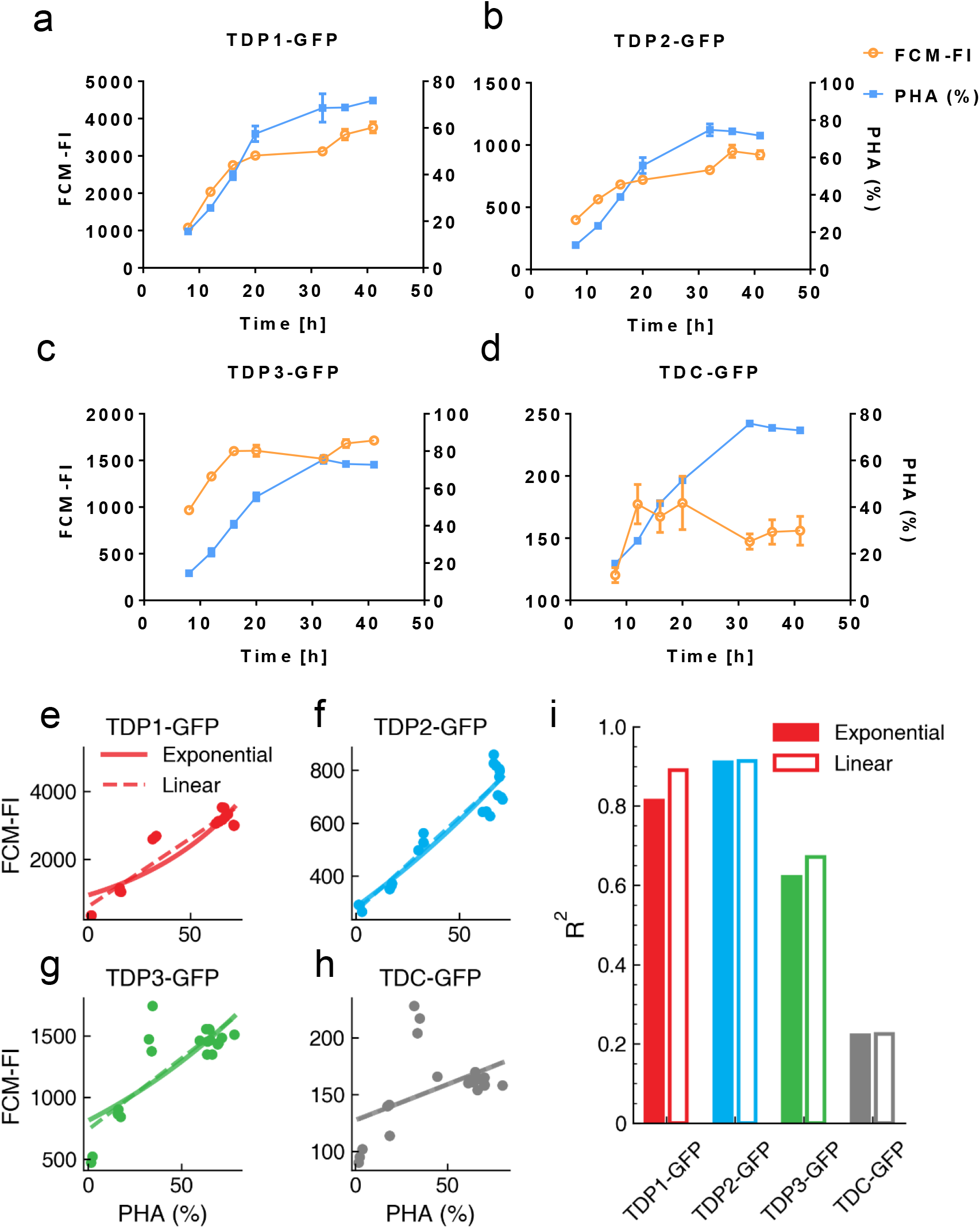
PHA contents related to fluorescence intensities (FI) generated by the three GFP labeled PhaPs in *H. bluephagenesis*, respectively. (**a-d**) Dynamic FCM-FI (orange hollow circle) related to PHA contents (blue solid square) generated by the cells of *H. bluephagenesis* TDP1-GFP, TDP2-GFP, TDP3-GFP and TDC-GFP grown under same conditions, respectively. (**e-h**) Exponential and linear functions were used to fit the relationship between the FCM-FI and the PHA contents generated by *H. bluephagenesis* TDP1-GFP, TDP2-GFP, TDP3-GFP and TDC-GFP grown under same conditions. (**i**) The R-square values of **Figs. 3e-3h** are shown in columns. The solid and hollow columns represent exponential and linear functions, respectively. All data represent the mean of n = 3 biologically independent samples and error bars show s.d.

### 3.4 The qPHA method with standard parameters generated from a microplate reader

Although the flow cytometry (FCM) can be used to quantify intracellular PHA contents as suggested above, the equipment is too heavy, large and expensive to be used for many labs. Low cost microplate readers are widely used equipment with possibility to replace flow cytometers. The difficulty there was to normalize the cell number by microplate readers in contrast of flow cytometers as they could measure the fluorescent intensity of single cells. The common normalization method depended on the OD_600_ is directly proportional to the cell numbers. But the relationship in the cell contains various content PHA is different. The intracellular PHA would influence OD600 value, resulted in misestimation of cell numbers (Martinez & Deziel, 2020).

A new method to quantify the PHA content in living cells was developed based on the OD_600_ and GFP fluorescence intensity (FI) obtained from the microplate reader. Firstly, OD_600_ (O)was reported to be in direct proportional to the mean PHA content (PHA) for all cells and the total cells number (n) in the well [Eq. (1)] (Koch, 1961; Martinez & Deziel, 2020; Stevenson et al., 2016). Secondly, the GFP fluorescence intensity (F) of each well, is linearly dependent on the mean fluorescence of all cells measured by FCM (f) mentioned previously [Eq. (2)] (Pasquier et al., 2013). The k1-3 and b1-3 were the adjustment parameters for the following equations.

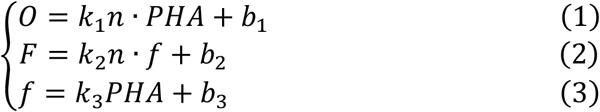

Since the linear function effectively fits the relationship between the FCM-FI and the PHA content as we discussed above, it can be described as Eq. (3). When solving Eqs (1)–(3), the PHA content can be calculated based on the OD_600_ and GFP fluorescence intensity (FI) as described in Eq. (4):

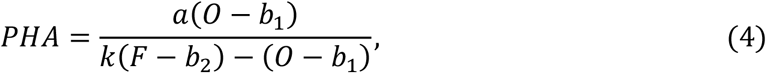

Where *a = (k_2_k_3_ + b_2_* and k = k_1_/(k_2_k_3_). Otherwise, the exponential function form of Eq. (3) generated different relationships among PHA (%), OD_600_ and FI, as described in Supplementary material.

For another part, the strain used in this experiment is based on *H. bluephagenesis* TDH4Δ*phaP1*, which was able to produce poly(3-hydroxybutyrate-*co*-4-hydroxybutyrate) or P(3HB-*co*-4HB) in form of large PHA granules in each cell. The T7-like promoter system and the *phaCAB* operon were inserted into the genome of *H. bluephagenesis* TDH4Δ*phaP1* to allow isopropyl-β-d-thiogalactoside (IPTG) induction of more PHA synthesis (Zhao et al., 2017). As *phaP1* was deleted in this strain for forming large PHA granules in each cell, *phaP2* remained as the choice to fuse with GFP for indicating the intracellular PHA content. Thus, the GFP labeled PhaP2 strain *H. bluephagenesis* TDH4Δ*phaP1*P2-GFP was constructed.

Firstly, PHA production by *H. bluephagenesis* TDH4*Δphap1*P2-GFP was studied under 1 g/L urea concentration and found to be highly related to the FCM-FI in both linear and exponential functions, respectively (Fig. 4d). These results are consistent with the above experiments. Growth of *H. bluephagenesis* TDH4*ΔphaP1*P2-GFP were repeated twice to compare their results in terms of OD_600_ and GFP fluorescence intensity (FI) using the microplate reader, with small deviations observed at late stage of growth (Figs. 4a-4c). Eq. (4) was employed to fit the data from the two above experiments, and calculated the R-squares, respectively (Fig. 4e). The parameters were calculated to be *a = 1.59 × 10^9^*, *b_1_ = −2.34*, *b_2_ = −7.60 × 10^2^* and k = 4.49 × 10^5^, which are defined as the standard parameters. The R-square values from both growth experiments were larger than 0.95, and the PHA content obtained from GC analysis (points) with the PHA content calculated by the curve of the Eq. (4) using the above parameters showed in the 3D graph and the time profile, respectively (Figs. 4e-4f). In addition, 2D projection graph showed the relationship between the OD_600_ and PHA content as well as FI and PHA content, respectively. (Figs. 4g-4h). All the above calculated PHA contents agreed with real values from GC analysis (Fig. 4). Eq. (S4) based on the exponential function form of Eq. (3) [Eq. (S3)] also could fit well with the two above experiment data (see Supplementary material). To simplify and standardize the qPHA method, the laborious computation procedure was encapsulated as a software that was described in Date S1 and could be download at GitHub (https://github.com/EggYouKnow/qPHA). With this software, users could get personalized parameters by importing series of FI, OD_600_ and PHA content experimental data. Also, the PHA content could be calculated based on FI and OD_600_ with fitted parameters in this software. The microplate-based method named qPHA was established.

**Figure. 4.**
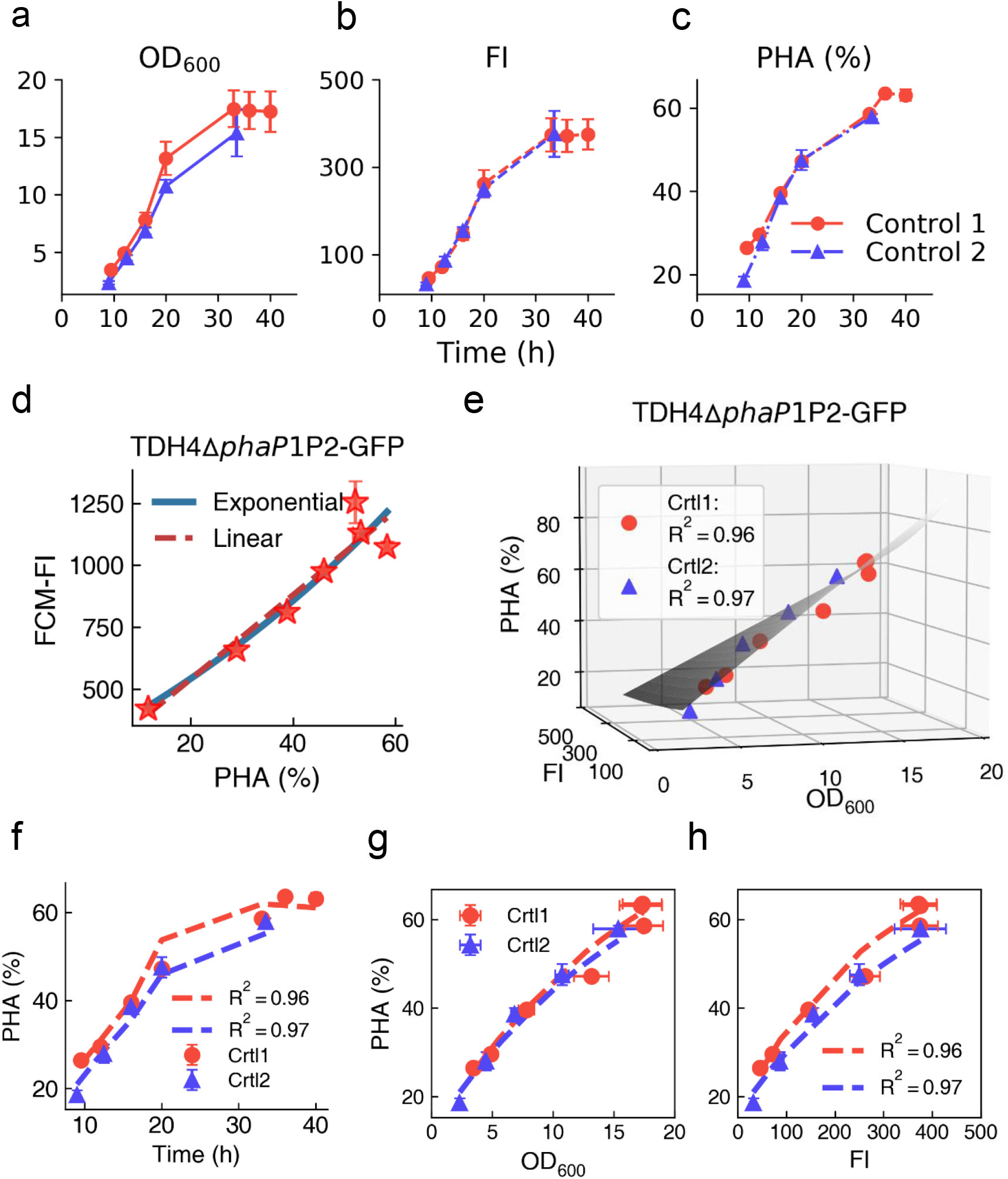
OD_600_ and fluorescence intensity (FI) related to PHA accumulations fluorescence intensity (FI) generated by *H. bluephagenesis* TDH4ΔphaP1P2-GFP grown in shake flasks. (**a-c**) fluorescence intensity, OD600 and PHA content from two independent shake flask studies using *H. bluephagenesis* TDH4*ΔphaP1*P2-GFP. (**d**) Relationships of FCM-FI and the PHA content fitted using linear (red dashed line) and exponential (blue line) functions, respectively. The stars represent experimental data (n=3 samples). (**e-h**) Relationship of PHA content, OD600 and FI fitted to Equation 4 and their standard parameters *a = 1.59 × 10^9^*, *b_1_ = −2.34*, *b_2_ = −7.60 × 10^2^*, *k = 4.49 × 10^5^* Fitting of Equation 4 (gray surface) to experimental data in a 3D version. (**f**) Pseudo-time series of fitting results (dashed line) compared to the experimental sequential data. (**g-h**) Fitting results in 2D projection graphs. (**a-c**) & (**e-h**) Red circles (Control 1 or Crtl1) and blue triangles (Control 2 or Crtl2) stand for the mean levels of data from two independent experiments. Error bars represent the standard deviation estimated with n=3 samples for each experiment.

### 3.5 Quantification via the qPHA method for PHB and P(3HB-*co*-4HB)

The qPHA method established to measure OD600 and GFP fluorescence intensity (FI) via microplate reader for PHA content, was further exploited for quantifications of PHB synthesis from different substrates. It was reported that a proper concentration of acetate (such as 6 g/L) could significantly enhance the intracellular PHA synthesis in *H. bluephagenesis* by consuming excess NADH (Ling et al., 2018). *H. bluephagenesis* TDH4Δ*phaP1*P2-GFP grown in the presence and absence of acetic acid were analyzed via the qPHA method to find well fit the curve generated from the standard parameters ( Fig. 5a). In the time sequence diagram, the FI calculated PHA contents agreed well with the experiment data, indicating that the qPHA method is suitable for PHA quantification in different substrates such as glucose and acetic acid (Fig. 5b).

**Figure. 5.**
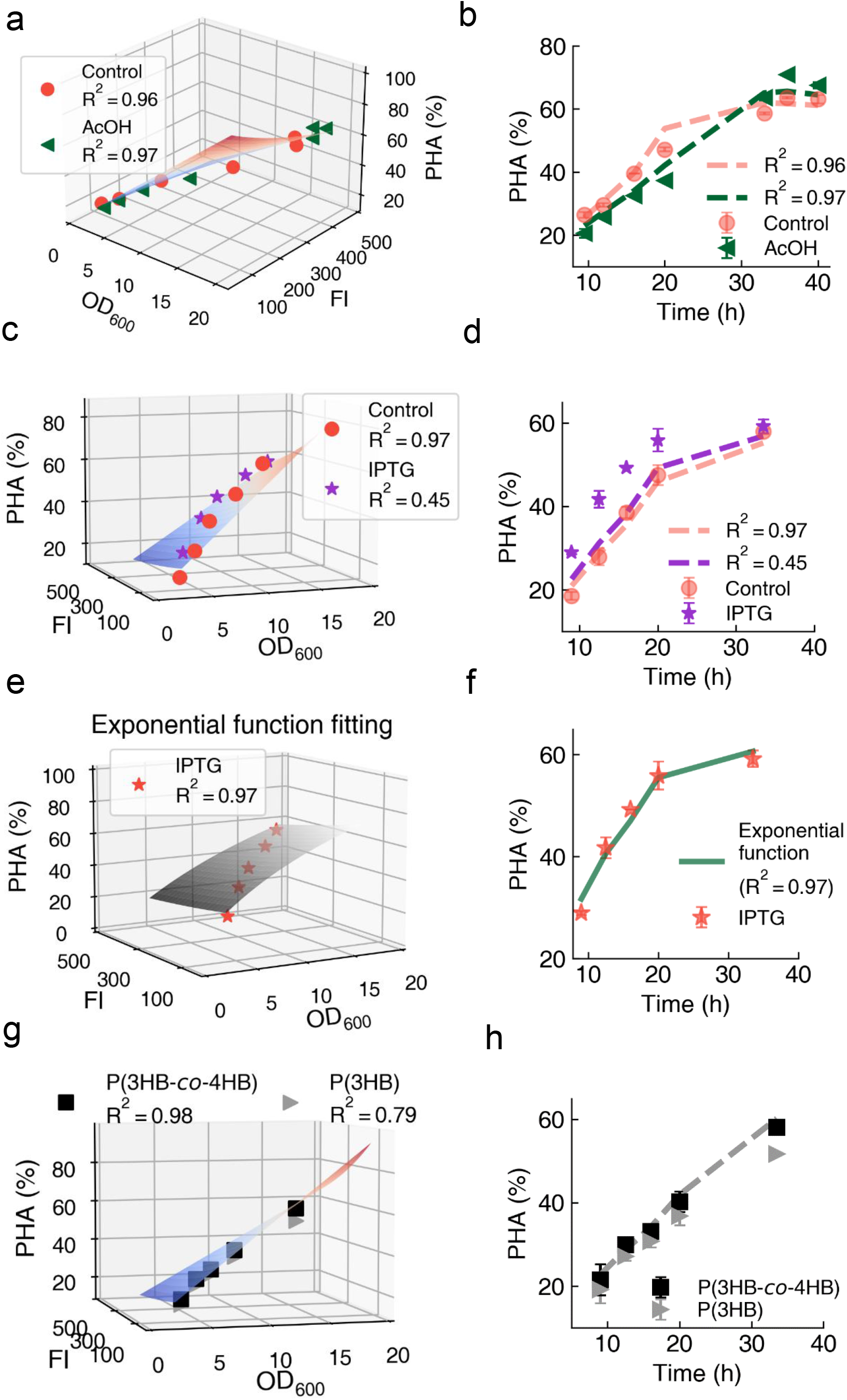
Applications of qPHA method under different PHB level and different PHA composition. (**a**) The adding acetic acid (AcOH) group and the control group were fitted by Equation 4 with the standard parameters which showed the relationship of the PHA content, OD600 and fluorescence intensity by 3D view with R-square values. Standard parameters: *a = 1.59 × 10^9^*, *b_1_ = −2.34*, *b_2_ = −7.60 × 10^2^*, *k = 4.49 × 10^5^*. (**b**) The experiment data and the fitting curve of Fig. 5a were showed in time sequence diagram. (**c**) The adding IPTG group and the control group were fitted by Equation 4 with the standard parameters showed the relationship of the PHA content, OD_600_ and fluorescence intensity by 3D view with R-square values. Standard parameters: *a = 1.59 × 10^9^*, *b_1_ = −2.34*, *b_2_ = −7.60 × 10^2^*, *k = 4.49 × 10^5^*. (**d**) The experiment data and the fitting curve of Fig. 5c were showed in time sequence diagram. (e) The exponential function (Supplementary Materials) was used and values of b1, b2 were fixed during parameter estimation. And the experiment data (red star) and the adjusted fitting surface were shown with R-square values. The adjusted parameters, *a’ = −1.81 × 10^−5^*, *k_4_ = 7.28 × 10^−4^*, and *c = − 1.11* (**f**) The experiment data and the fitting curve of exponential function with new estimated parameters were showed in time sequence diagram. (**g**) Calculated P(3HB-*co*-4HB) group and the group calculated P(3HB) group alone were fitted by Equation 4 with the standard parameters which showed the relationship of the PHA content, OD600 and fluorescence intensity by 3D view with R-square values. Standard parameters: *a = 1.59 × 10^9^*, *b_1_ = −2.34*, *b_2_ = −7.60 × 10^2^*, *k = 4.49 × 10^5^*. (**h**) The experiment data and the fitting curve of Fig. 5g were showed in time sequence diagram. All data represent the mean of n = 3 biologically independent samples and error bars show s.d.

In another study, IPTG was used to induce the expression of additional PHB synthesis operon *phaCAB* of *C. nector* for enhanced PHA accumulation by *H. bluephagenesis* TDH4Δ*phaP1*P2-GFP harboring both MmP1 RNA polymerase encoding gene and the P_MmP1-LacO_-*phaCAB* module (Shen et al., 2019; Zhao et al., 2017). Under this situation, the qPHA method with established standard parameters became unsuitable with low R-square (Figs. 5c-5d). Parameters of the functions (1) and (2) are related to the equipment, which did not change in the IPTG experiment. The parameters of function (3) have been changed for the heterologous expression of *phaCAB*. Moreover, the adjustments of k3 and b3, that are the paraments of Eq. (3), could not significantly improve the fitting results generated from the IPTG induction. As a result, the relationship of FCM-FI and PHA content under *phaCAB* overexpression condition was considered as the exponential function. When an exponential function form of Eq. (3) [Eq. (S3)] was used to replace the linear function [Eq. (3)], the PHA content calculated by FI and OD_600_ fit well with the experimental data of PHA content with a R-square over 0.95 (Figs. 5e and 5f). Detailed parameters acquisition was listed in Method.

To study if the method is also applicable to other PHA copolymers, such as P(3HB-*co*-4HB) containing various amounts of 4-hydroxybutyrate (4HB) with adjustable elongation at break (Chanprateep et al., 2010; Chen et al., 2017; Cong et al., 2008), *H. bluephagenesis* TDH4Δ*phaP1* containing 4HB-CoA transferase in genome was studied as it is able to produce P(3HB-*co*-4HB) in the presence of γ-butyrolactone in the culture medium(Shen et al., 2019). When grown in an optimal 5 g/L γ-butyrolactone in the MM60 medium (Shen et al., 2018), P(3HB-*co*-4HB) contents fit the FI values and OD600 well with the R-square of 0.98 (Fig. 5g). In contrast, PHB content fitting to the FI value and OD_600_ generated a R-square of only 0.79 (Fig. 5g). These results are more clearly in the time profile (Fig.5h). In conclusion, this qPHA method can be adjusted for quantifying not only homopolymer PHB but also copolymer P(3HB-*co*-4HB).

### 3.6 The qPHA method to monitor PHA synthesis in bioreactors

It has been impossible to conduct rapid PHA synthesis quantification for process monitoring and control due to the time and labor consuming GC analysis process, while this is important to adjust the processes for enhanced PHA production. To achieve *in situ* monitoring intracellular PHA contents, the qPHA method was employed to follow the PHA synthesis by *H. bluephagenesis* TDH4Δ*phaP1*P2-GFP incubated in a 7.5 L bioreactor (Fig. 6). Small amounts of culture fluids (mL) were taken during the fermentation process lasting about 48 h, respectively (Fig. 6b), diluted to an OD600 less than 0.5 before conducting the FI analysis using a fluoro-spectrophotometer (Tan et al., 2011). As fluoro-spectrophotometers are less costly than microplate readers and flow cytometers, they are more suitable for rapid quantification during industrial PHA productions. The PHA contents were generated by FI and OD_600_ in minutes and agreed very well with the GC results with a R-square of 0.99 (Figs. 6a-6d).

**Figure. 6.**
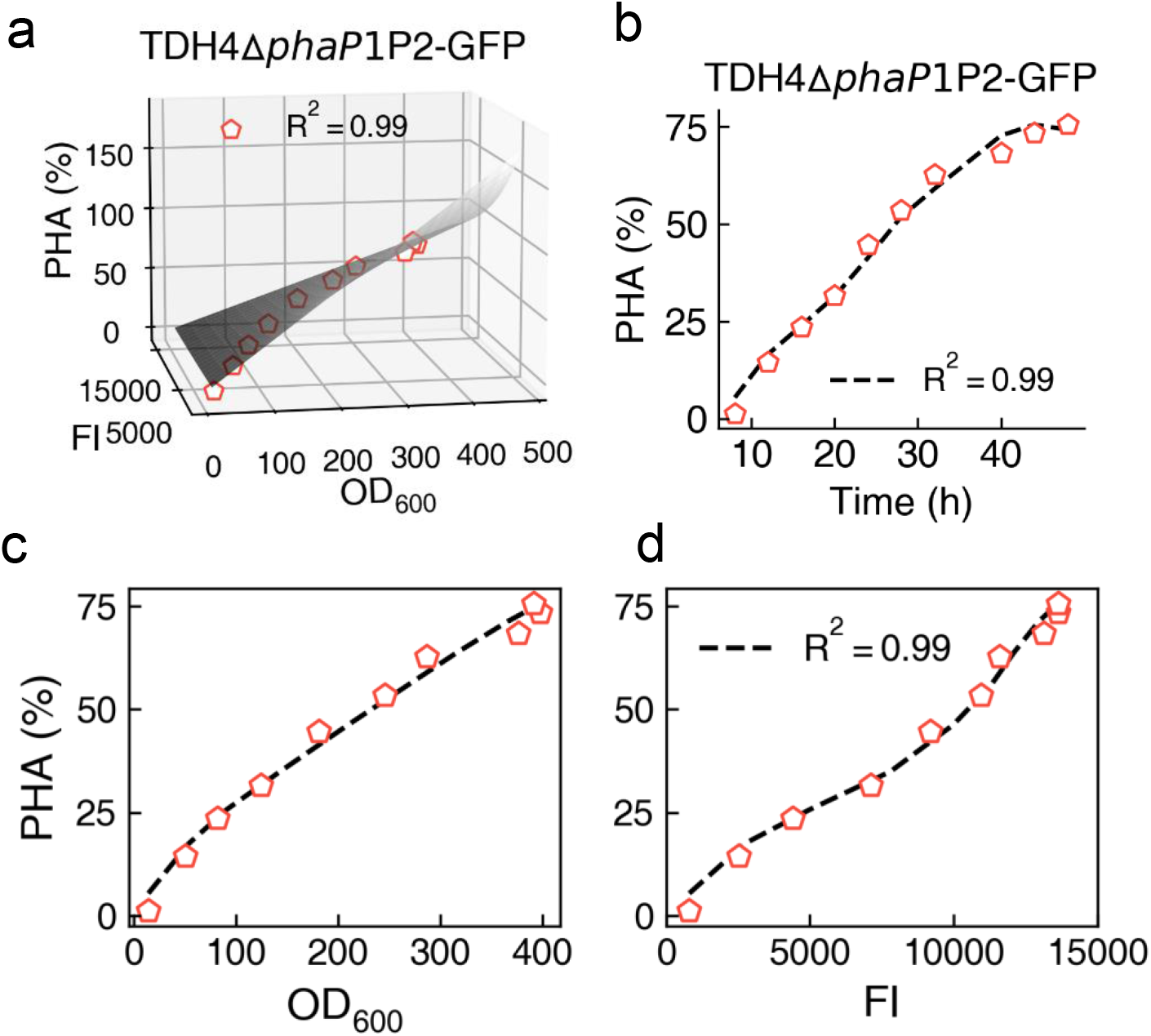
*In situ* quantitation application of the qPHA method for evaluation of intracellular PHA accumulations by. *H. bluephagenesis* TDH4Δ phaP1 P2-GFP grown in 7.5 L fermenters. **(a)** Fitting of Equation 4 (gray surface) to experimental data in a 3D version. (**b**) Pseudo-time series of fitting results (dashed line) compared to the experimental sequential data. **(c-d)** Fitting results in 2D projection graphs. (**a-d**) Red hollow pentagon points represent the PHA contents of the cells incubated in the fermenter.

Because of the use of the fluoro-spectrophotometer and the changing culture conditions, the parameters for Equations 4 were updated with new given values of parameters *a = 1.01×10^8^, b1 = 1.33×10^1^*, *b2 = −6.88 × 10^3^, k = 2.54 × 10^4^)*, the R-square reached 0.99 indicating satisfactory agreements between FI, OD600 and PHA contents (Figs. 6a-6d). Results demonstrated the potentials to apply the qPHA method for rapid and convenient fermentation process monitoring and controls.

### 3.7 Discussion

Polyhydroxyalkanoates have been considered as a promising bioplastic that can be synthesized and processed into commodity bioplastics or high value medical materials (Chen & Jiang, 2018; Moradali & Rehm, 2020). For the process control and monitoring during the PHA synthesis, lower-cost and less time-consuming quantification of intracellular PHA is very significant. Therefore, for the first time, the GFP labeled native phasin was constructed as an indicator to analyze the intracellular PHA content (Fig. 1). When grown in 1 g/L urea, cells were grown with sufficient PHA production meeting the positive correlation to FCM-FI (Fig. 2). Both PhaP1 and PhaP2 were chosen as suitable targets to fuse with GFP for indicating the intracellular PHA content among the three PhaPs in *H. bluephagenesis* (Fig. 3). Moreover, to simplify the GFP PHA quantification applications, the method was successfully extended from flow cytometer to lower-cost microplate reader. Based on the three equations, the intracellular PHA content can be modeled with FI and OD_600_ that can be both measured via a microplate reader using standard parameters (Fig. 4). To simplify the usage, all the calculated processes can be conducted using software list in Supplementary Materials.

As for the qPHA method applications during fermentations, the substrate- and gene-expression-level adjustments are two common ways to regulate the production of bioproducts (Jiang et al., 2011; Zhao et al., 2017). The qPHA method combined with the standard parameters, was found suitable for intracellular PHA quantification in the presence of acetic acid in the culture medium (Figs. 5a-5b). Under conditions of disturbance for heterologous gene expression of the PHA synthesis pathway, the exponential function [Eq. (S3) in Supplementary Materials] was used to describe the relationship between FCM-FI and PHA content, the experiment data fit well with the calculated results with an R-square over 0.95 (Figs. 5e-5f). The parameters in the above equations can be easily adjusted to meet various experimental conditions. The relationship of FCM-FI and PHA content can be updated and reconstructed when PHA production ability was changed, as in the case of over expressing *phaCAB* operon. The qPHA method was also applied to quantify intracellular copolymer PHA such as P(3HB- *co*-4HB) with significantly fitting results (Figs. 5g-5h). Furthermore, low cost spectrophotometer was used *in situ* to successfully monitor PHA industrial production during the fermentation process (Fig. 6). Importantly, the method detects PHA production in living cells instead of dead ones, which provides the observation of PHA real-time formation in microfluidics under microscope (see Supplementary material)).

The qPHA method can rapidly provide the intracellular PHA content with several microliters sample in minutes compared to the traditional GC analysis method or the fluorescence staining method. For the GC analysis method, the whole processes from sample to data require nearly two days and need at least 20 mg dry sample. For fluorescence staining methods, it requires time for incubation and the staining will lead to cell death or even false results (Lee et al., 2013). With the shortest analysis time, the qPHA method can be conveniently applied to monitoring the bioreactor fermentation, contributing to real-time fermentation control. At the same time, the qPHA method does not influence cell growth, indicating the possibility that the sampled cells be reinoculated after the intracellular PHA content analysis (see Supplementary material)). FCM sorting for single cells could be conducted for higher PHA producing cells in future experiments. Otherwise, the PHA yield in 1.0 g/L urea concentration group are even a little higher than the 0.5 g/L group, indicating the PHA productivity is not hampered in qPHA method, which related with specific nitrogen concentration (see Supplementary material).

As the granules associated proteins for enveloping PHA, the amphiphilic protein PhaPs are commonly found in native PHA production strains including *Aeromonas hydrophila*, *C. necator* and even archaea *Haloferax mediterranei* or cyanobacteria *Synechocystis sp. strain PCC 6803*(Cai et al., 2012; Hauf et al., 2015; Zhao et al., 2016). It was reported that the PhaP1*Cn* is also in direct proportional to PHA quantity in *C. necator* (York et al., 2001a; York et al., 2001b). The qPHA method can be applied in other PHA producers mentioned above in which PhaPs are under the control of PHA formation.

Fluorescence proteins as visible probes possess advantages for observing biological processes. Here, fusion fluorescence protein was developed for the product *in situ* quantity assays, the fusion method provides possibility for quantification on other bio-products correlated to certain protein-levels, quantifying that proteins are equal to quantitation of the metabolites.

## 4. Conclusion

In conclusion, the qPHA method has been successfully establish to quickly quantify intracellular PHA contents in living cells. The method should be able to promote the development of the PHA industry which is limited by process development due to the lack of process monitoring means.

E-supplementary data of this word can be found in online version of the paper.

## Acknowledgements

We thank Prof Victor de Lorenzo for donating the SEVA series vectors.

## Funding

Ministry of Science and Technology of China (Grant No. 2018YFA0900200) (**Guo-Qiang Chen**)

National Natural Science Foundation of China (Grant No. 31870859; No. 21761132013; No. 31961133019; NO. 31961133017; NO. 31961133018) (**Guo-Qiang Chen**)

MIX-UP, European Union’s Horizon 2020 research and innovation program (No. 870294) (**Guo-Qiang Chen)**

## Author contributions

Conceptualization: Fangting Li, Guo-Qiang Chen, Xu Liu, Dianjie Li

Methodology: Xu Liu, Zonghao Zhang, Shuang Zheng, Jingpeng Zhang Investigation: Xu Yan

Visualization: Xu Liu, Dianjie Li

Supervision: Fangting Li, Guo-Qiang Chen, Fuqing Wu Writing—original draft: Xu Liu, Dianjie Li Writing—review & editing: Guo-Qiang Chen

## Competing interests

All other authors declare they have no competing interests.

## Data and materials availability

All data are available in the main text or the Supplementary materials.

## Notes

### Competing Interest Statement

The authors have declared no competing interest.

